# Deep learning of representations for transcriptomics-based phenotype prediction

**DOI:** 10.1101/574723

**Authors:** Aaron M. Smith, Jonathan R. Walsh, John Long, Craig B. Davis, Peter Henstock, Martin R. Hodge, Mateusz Maciejewski, Xinmeng Jasmine Mu, Stephen Ra, Shanrong Zhao, Daniel Ziemek, Charles K. Fisher

## Abstract

The ability to predict health outcomes from gene expression would catalyze a revolution in molecular diagnostics. This task is complicated because expression data are high dimensional whereas each experiment is usually small (e.g., *∼*20,000 genes may be measured for *∼*100 subjects). However, thousands of transcriptomics experiments with hundreds of thousands of samples are available in public repositories. Can representation learning techniques leverage these public data to improve predictive performance on other tasks? Here, we report a comprehensive analysis using different gene sets, normalization schemes, and machine learning methods on a set of 24 binary and multiclass prediction problems and 26 survival analysis tasks. Methods that combine large numbers of genes outperformed single gene methods, but neither unsupervised nor semi-supervised representation learning techniques yielded consistent improvements in out-of-sample performance across datasets. Our findings suggest that using *l*_2_-regularized regression methods applied to centered log-ratio transformed transcript abundances provide the best predictive analyses.

## I. INTRODUCTION

The potential to tailor therapies for individual patients rests on the ability to accurately diagnose disease and predict outcomes under various treatment conditions. Predictors based on high-throughput ‘omics technologies hold great promise, but a number of technical challenges have limited their applicability [1]. Phenotypes may be complex—involving contributions from large numbers of genes—but ‘omics data are so high-dimensional that exploring all possible interactions is intractable. This situation is further complicated by the small sample sizes of typical biological studies and by large systematic sources of variation between experiments [2],[3]. However, recent developments in machine learning have raised hopes that new computational methods integrating data from many studies may be able to overcome these difficulties. Accurate prediction of phenotype or endpoint(s) from ‘omics data would usher in an era of molecular diagnostics [4],[5].

Machine learning methods often benefit from large datasets where learning complex relationships is feasible. Although individual biological experiments tend to be small, relatively large amounts of ‘omics data are available in public repositories. For example, hundreds of thousands of samples from human RNA sequencing (RNA-seq) experiments are available from the recount2 and ARCHS4 databases [6–8]. Still, these data cover a wide variety of tissues and diseases. Moreover, there are no specific diseases with large numbers of samples and, in many cases, the metadata are not sufficient to determine basic experimental facts like the tissue of origin [9]. As a result, leveraging these data to improve prediction tasks will require machine learning techniques that can learn from large, heterogeneous datasets.

Genes rarely act in isolation, so it is reasonable to expect that combinations of genes may be more effective than individual genes for predicting phenotypes. For example, linear models operating on RNA-seq data create predictors from a weighted combination of gene expression values. However, some of these features could reflect biological processes that are involved in multiple phenotypes. Many previous analyses have explored this possibility by creating complex features that incorporate biological knowledge from gene sets [10, 11], ontologies [12], or interaction graphs [13–15]. More recently, machine learning methods used with principal components analysis [16], autoencoders [17–20], or other neural network architectures have been developed to discover such features by analyzing large transcriptomics datasets. If these learned features capture biologically relevant processes, then predictive models built from those features should outperform models built directly from relative transcript abundances.

In this work, we present a comprehensive analysis of phenotype prediction from transcriptomics data with a particular emphasis on representation learning. Using the recount2 database [7], we systematically explored the impact of normalization techniques, gene sets, learned representations, and machine learning methods on predictive performance for a set of 24 binary and multiclass prediction problems and 26 survival analysis tasks. In total, we analyzed thousands of predictive models using 5-fold nested cross validation to rigorously assess out-of-sample performance. We found that predictors that combined multiple genes outperformed single gene predictors, that logarithmic transformations outperformed untransformed relative expression measurements, and that for survival analyses larger gene sets outperformed smaller gene sets. However, neither unsupervised nor semi-supervised representation learning techniques yielded consistent improvement on out-of-sample predictive performance across datasets. In fact, *l*_2_-regularized regression methods applied directly to the centered log-ratio transform of transcript abundances performed consistently well relative to the other methods. Therefore we recommend treating that particular combination as a baseline method for predictive analysis on RNAseq data. Throughout this text we refer to the combination of *l*_2_-regularized regression methods applied directly to the centered log-ratio transform of transcript abundances as *the recommended model*.

## II. RESULTS

### Approach

A high-level description of our quality control, data processing, and machine learning analyses is provided in Figure 1. Details of these steps are provided in the online Methods.

**FIG. 1:**
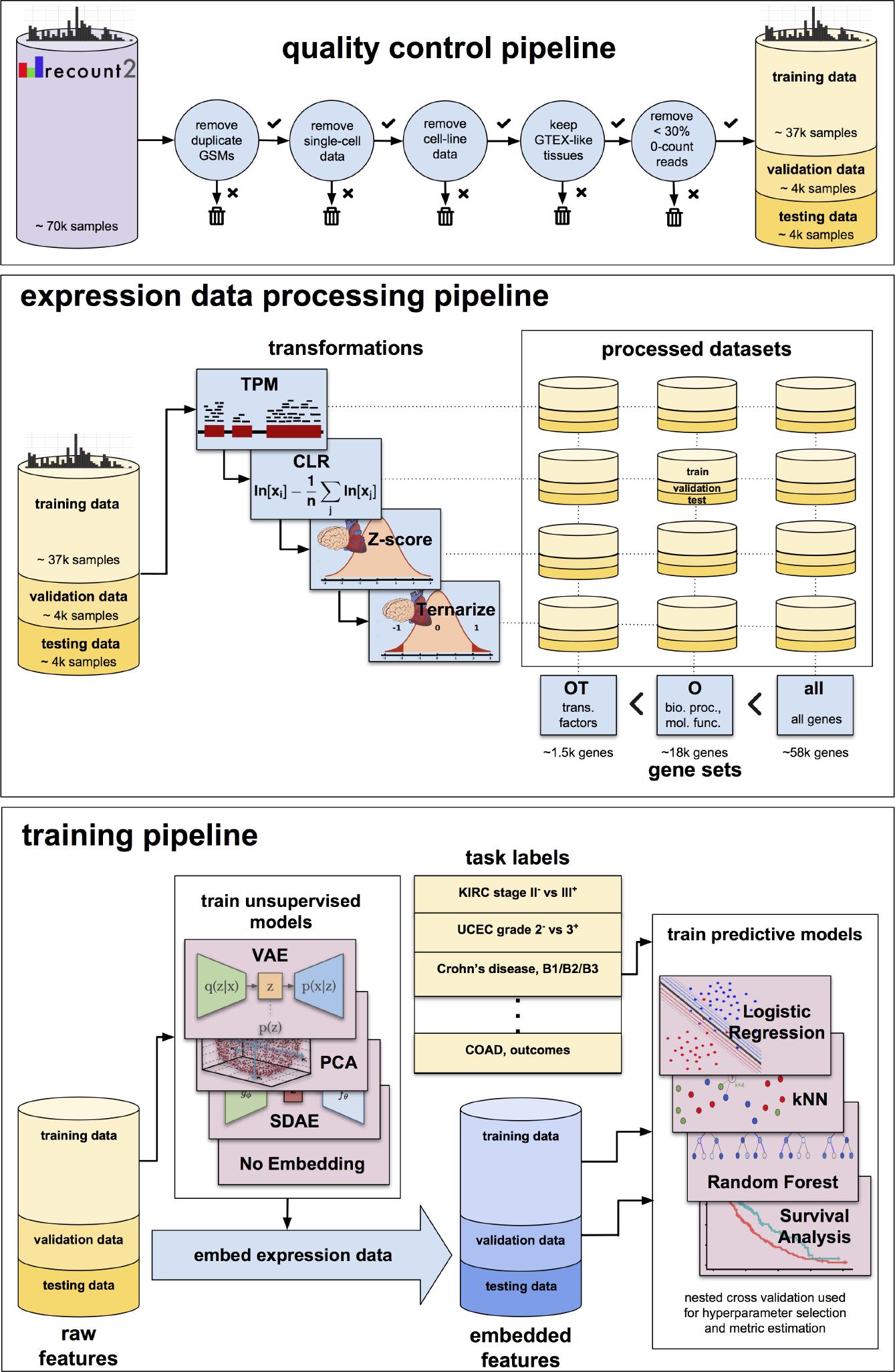
Schematic. An overview of the quality control, data processing, and training pipelines. Data from recount undergoes several quality checks at the sample and study level, resulting in a dataset of approximately 45,000 samples divided into training, testing, and validation datasets. Twelve different datasets are created from these data, each with a different gene set (**all**, comprising all genes; **O**, comprising key GO categories; **OT**, comprising **O** genes that are known transcription factors) and transform (“TPM”, transcripts per million; “CLR”, a centered-log-ratio transform of TPM; “Z-score”, a Z-score normalization of the CLR data relative to healthy tissue expression levels in GTEx; “Z-ternary”, a ternarization of Z-score). The training data is used to train unsupervised models capable of embedding the data (a “no embedding” model is also included, which does not alter the data). These embedded features, along with labels for individual tasks, are used to train a variety of supervised models. The supervised models are trained and evaluated using nested cross validation.

Briefly, we selected a subset of experiments from the recount2 database [7] that did not have sparse gene expression data and could be mapped to the same set of tissues covered in the Genotype-Tissue Expression (GTEx) project [21]. We assigned the various experiments to “training” (*∼*37k samples), “validation” (*∼*4k samples), and “test” sets (*∼*4k samples). All samples lacking suitable metadata for supervised learning were allocated to the training set. From metadata provided with recount2, the Gene Expression Omnibus [22], and The Cancer Genome Atlas (TCGA) Pan-Cancer Clinical Data Resource [23] we derived labels for 24 binary and multiclass and 26 survival analysis tasks. Descriptions of these tasks and their assignment to the training, validation, and test sets are provided in the online Methods.

We considered four different normalization methods to correct for variance introduced in the data collection and measurement process. We first converted the samples from counts to Transcripts Per Million (TPM) [24], a normalization which estimates relative molar concentration of transcripts in a sample. Under the operating assumption that transcript abundance is determinant of downstream biological function, TPMs should be the baseline quantification to work with from RNAseq. In contrast, raw counts (or CPMs) contain irrelevant counting bias stemming from variable transcript length. Likewise, the common alternative of FPKMs do not coherently measure relative molar concentration, because they rely on a sample-dependent normalization factor. As such, FPKMs are not a useful measure when processing samples which are not entirely technical replicates of a single tissue sample [24, 25]. Secondly we applied the centered log-ratio transformation (CLR) [26] to the TPM data to address the fact that RNA-seq data quantify relative, rather than absolute, gene expression [27],[28]. Since these two normalization methods do not account for the tissue of origin of the sample, we evaluated additional normalization methods based on differential expression with respect to normal tissue. The third normalization method converted the CLR transformed expression data from each sample to a tissue-normalized Z-score by subtracting the mean and dividing by the standard deviation of the associated tissue in GTEx. This mean and standard deviation of the CLR transformed expression data were computed across the GTEx data in recount2 for each annotated tissue type. Finally, a fourth ternarized normalization discretized the Z-scores into down-regulated (Z *< −*2), normal (*−*2 *<* Z *<* 2), or up-regulated (Z *>* 2) categories.

For each of these normalization approaches, we also explored three gene sets corresponding to transcription factors [29] (denoted **OT**), protein coding genes annotated as with biological processes or molecular functions in the Gene Ontology 12 (denoted **O**), and all genes provided by recount2 (denoted **all**). The **O** and **OT** gene sets are substantially smaller than the **all** gene set and allow exploration of the dependence on the number of genes. In total, we examined twelve different normalization-gene set combinations for each predictive problem.

We considered four different types of representations of the gene expression data learned by unsupervised models. First, supervised models were trained directly on the normalized expression data without a learned embedding. We also considered representations constructed with Principal Components Analysis (PCA), a Stacked Denoising Autoencoder (SDAE), and a Variational Autoencoder (VAE) trained on the 37k samples in the training set without any supervising information.

For each binary or multiclass prediction task, we trained a k-Nearest Neighbor (kNN) classifier, a Random Forest (RF), and an *l*_2_-regularized multinomial Logistic Regression (LR) on the normalized and transformed data using 5-fold nested cross validation. Using nested cross validation (Methods) is important because it accounts for performance variance that results from different hyperparameter choices (e.g., the number of nearest neighbors, the depth of the trees in the forest, or the strength of the regularization coefficient). An *l*_2_-regularized Cox proportional hazards model was used for all survival tasks, also with 5-fold nested cross validation. Binary tasks were compared using the Area Under the receiver operating characteristic Curve (AUC); multiclass tasks were compared using the accuracy, and survival tasks were compared using the concordance-index (C-index) [30],[31].

Our systematic model search covered four normalization methods, three gene sets, four representations, and three supervised algorithms totaling 144 comparison models for each of the 24 binary and multiclass tasks. For the survival tasks we used the same normalization methods, gene sets, and representations, but considered only one supervised algorithm (Cox proportional hazards). For comparison, we also trained linear predictors using the recommended method that were only allowed to use a single gene. The choice of gene was treated as a hyperparameter and optimized using 5-fold nested cross validation.

### Analyses

The predictive performance assessed through 5-fold nested cross validation varied considerably across and within the predictive problems (see Figure 2). Gene expression data improved predictive performance relative to random guessing in almost all cases, indicating that RNA-seq data do contain information that is broadly useful for out-of-sample prediction. Moreover, linear predictors that used the expression data from all genes generally outperformed models that only used a single, most predictive gene. It is still common to analyze genes independently in differential expression and regression analyses; our results indicate, however, that linear combinations of genes are significantly more predictive than individual genes. Although there was sizable variance in performance across tasks, predictive performance was not correlated with any obvious dataset characteristics such as the number of subjects.

**FIG. 2:**
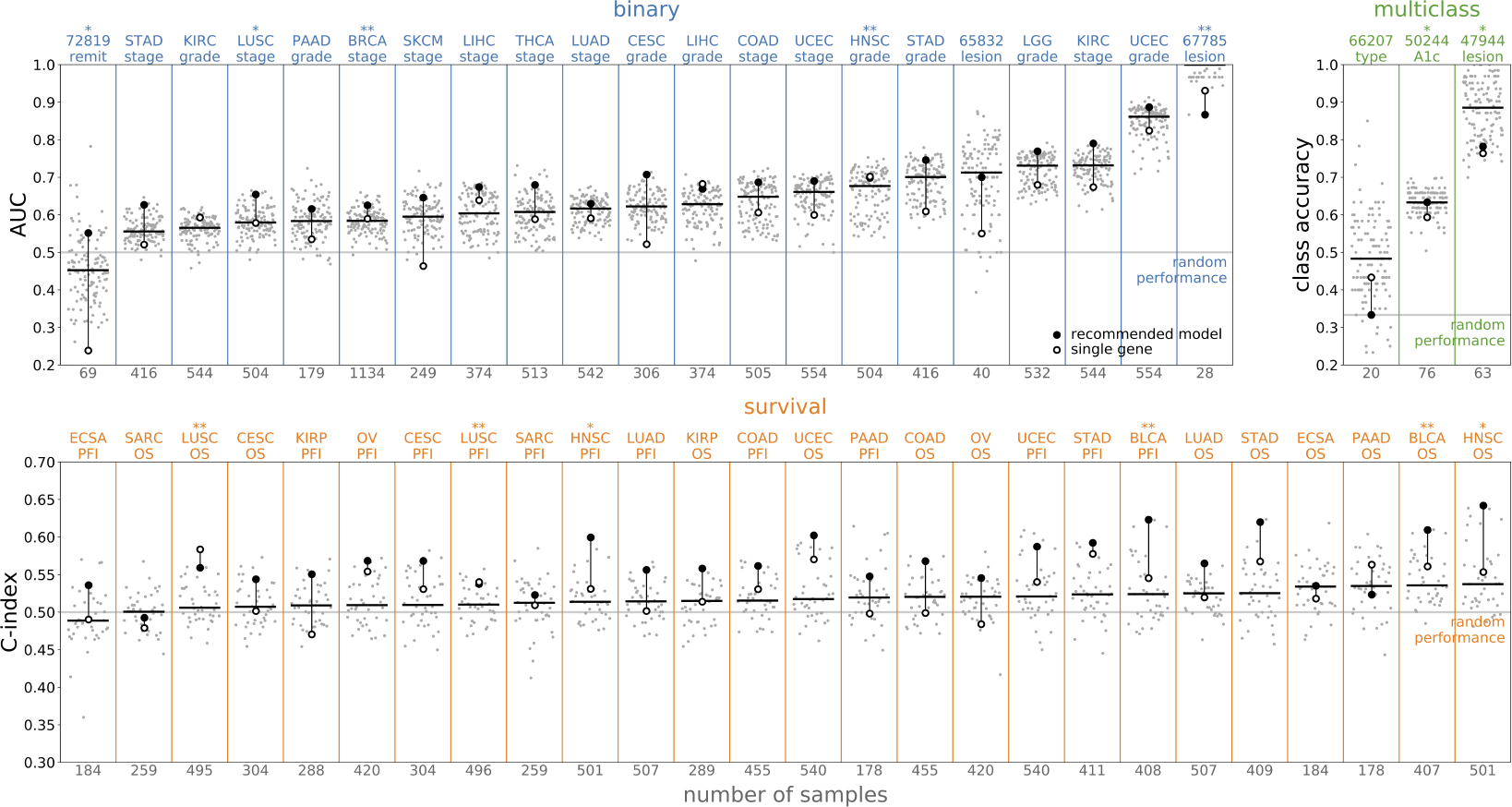
Performance by predictive task. The performance of all models on each task, ordered by the median performance on each task. The tasks are divided into three groups based on the type of label; the top row shows classification tasks (binary and multiclass) while the bottom shows survival tasks. Each task is labeled by an abbreviation at the top of the plot and the number of samples at the bottom; see the appendix for more details on each task. The task label has one star if the data is in the validation group and two stars if the data is in the test group. For each task, the gray points show the results over the entire set of models and the horizontal line shows their median. The filled black circle shows the performance of the recommended model, while the open black circle shows the performance of the best single gene model. The recommended model uses no embedding, **all** genes, and the CLR transform; the supervised model is logistic regression for the classifier tasks and a Cox proportional hazards model for the survival tasks. The recommended model is often among the best models on a problem and frequently outperforms the best single gene model; the primary exception is the pancreatic adenocarcinoma overall survival (PAAD OS) dataset.

In order to compare the effects of the gene set size and transformation, it is helpful remove between-task variance and then to aggregate results across tasks. To remove the between-task variance, we defined a shifted statistic in which we subtracted the median value of all models on the same task. For example, the AUC for the random forest classifier on the STAD stage dataset was shifted by subtracting the median AUC for all of the binary classifiers trained on the STAD stage dataset. Averages of the shifted statistics across predictive problems can be easily interpreted: if the value is less than zero then the method underperformed the median, whereas the method outperformed the median if the value is greater than zero.

Within-task variance in predictive performance was partially explained by the choice of gene set and normalization method (see Figure 3). Because the number of samples in each dataset was much smaller than the number of genes annotated in recount2, we hypothesized that using prior knowledge to select small, biologically relevant gene sets based on the Gene Ontology or transcription factor activity would improve out-of-sample predictive performance by preventing overfitting. However, this hypothesis was not supported by our analyses. The choice of gene set made no difference for the classification problems, whereas the smaller gene sets underperformed on the survival tasks. The log-transformed normalization methods slightly outperformed TPMs, and the Z-score normalization performed the best, on average. Performance improvements of Z-score normalization relative to CLR were small, however, and we do not think that the small gains justify the additional complexity introduced by referencing each sample to an external dataset (i.e., GTEx).

**FIG. 3:**
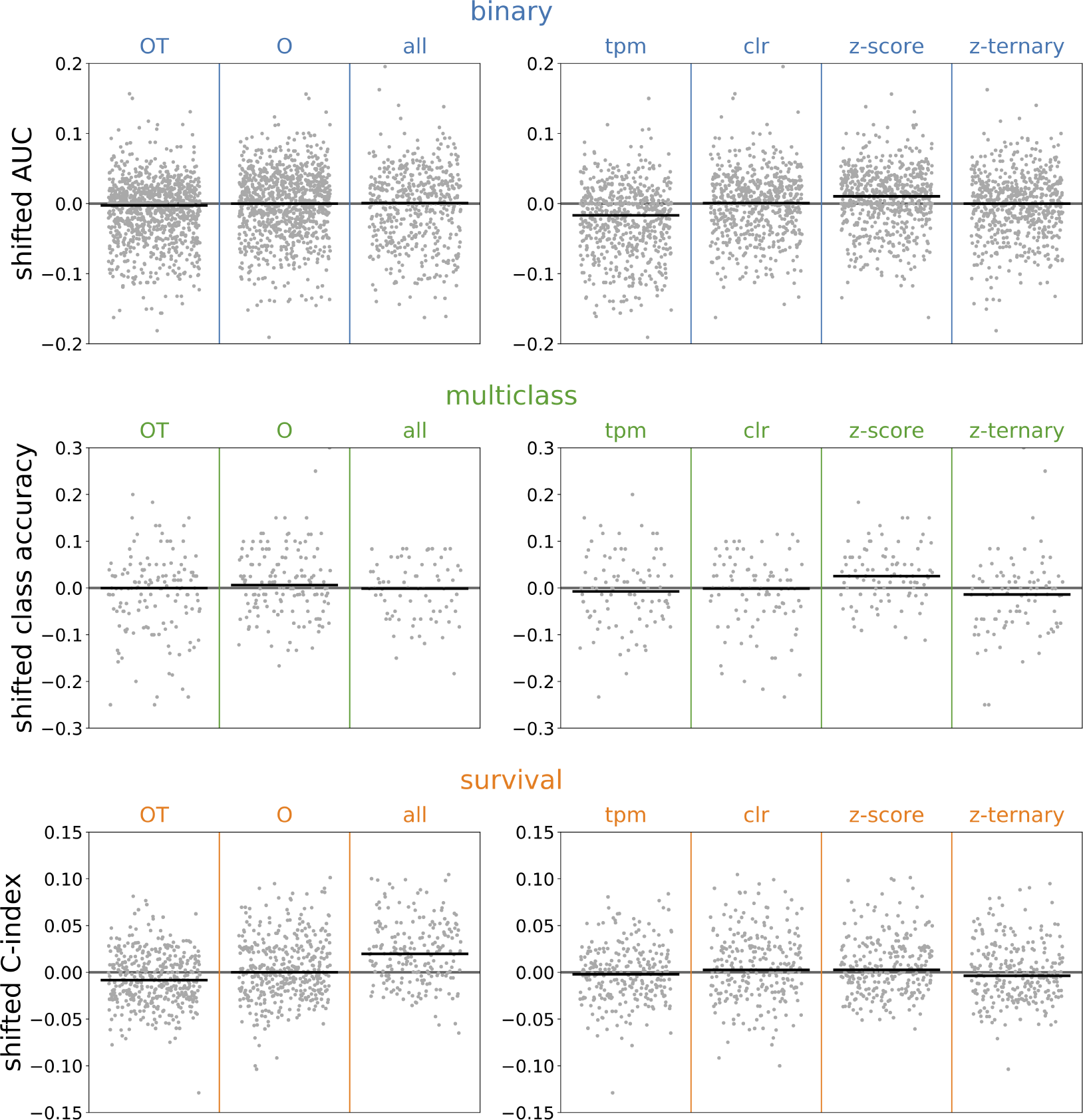
Performance by gene set and normalization. The performance of all models on each gene set (left column) and transform (right column). The results are divided by row into binary, multiclass, and survival tasks. For each gene set or normalization and task type, the gray points show the shifted statistics computed by subtracting the median of all models trained on the same task as a given model, and the black line is the median taken across all models and tasks.

Next, we examined differences in absolute performance between the kNN, RF, and LR models on the classification problems (only a linear Cox proportional hazards model was tested on the survival tasks). As shown in Figure 4, the kNN classifier consistently underperformed the RF and LR classifiers. The RF was the best performing method for thirteen tasks, LR for nine tasks, and kNN for two tasks, but LR was more consistent than RF and had better average performance.

**FIG. 4:**
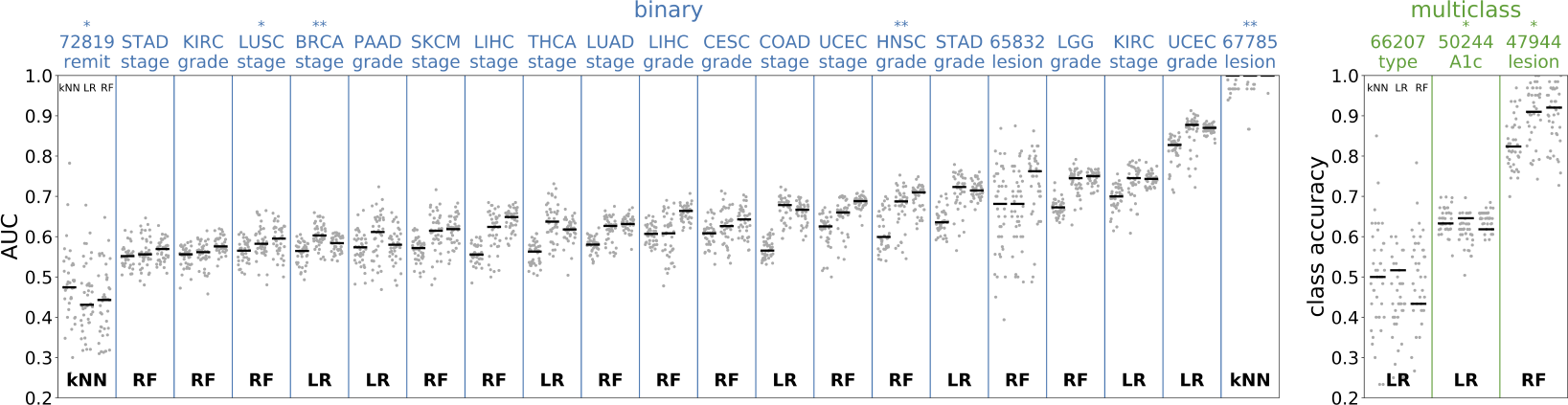
Performance by supervised model. The classifier tasks are shown in the same format as Figure 2. In this figure, the model results are divided into groups based on which supervised model is used, kNN (k-nearest neighbors), LR (logistic regression), or RF (random forest). The three horizontal lines for each task show the median result for each of these supervised models in this order (shown in the upper left of the first column plot). For each task, the supervised model with the best median result is shown at the bottom of the plot. While the median over the RF results is most frequently best (for thirteen tasks, compared to nine for logistic regression and two for *k*-nearest neighbors), the best performing logistic regression models are more consistently high performing among models.

Gene expression data are very high dimensional, with the number of genes ranging from *∼*1.5k in the transcription factor gene set to *∼*56k in the gene set consisting of all genes annotated in recount2. In contrast, the supervised task datasets typically consisted of only a few hundred samples. Moreover, it seems unlikely that genes actually coordinate in a linear fashion to generate complex phenotypes. Therefore we hypothesized that predictive performance could be improved by training predictors on lower dimensional representations derived from unsupervised analyses of the *∼*37k unlabeled samples in the training set. One could also view these analyses as a type of transfer learning, in which biological knowledge derived from the analysis of one dataset is used to inform the analyses of another.

The first feature representation that we considered was a Principal Components Analysis (PCA) with 512 latent dimensions. These principal components are orthogonal linear combinations of expression values that represent the directions of largest variance in the training set. Together, the 512 principal components we used explained the majority of the variation in the transcriptional datasets (Figure 9). We found that using PCA derived representations as features decreased the out-of-sample performance of downstream predictive analyses (Figure 5). Therefore, we do not recommend using features derived from PCA of large RNA-seq compendia for predictive analyses.

**FIG. 5:**
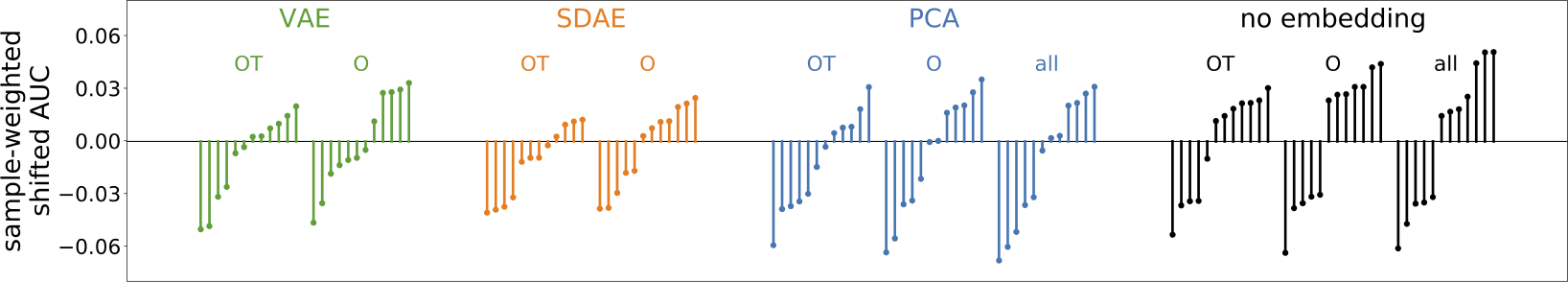
Performance by unsupervised model and gene set. The binary task performance of each unique model type is shown, grouped by unsupervised model and gene set. A model type is a combination of unsupervised model, supervised model, gene set, and normalization; for example, the recommended model is one model type. Each model type is a single line on this plot. The performance shown is the average of shifted AUCs across binary tasks, weighted by the number of samples in each task to reduce the effect of fluctuations in tasks with fewer samples. There are four unsupervised model types, VAE (variational autoencoder), SDAE (autoencoder), PCA (principal components analysis), and no-embedding (in which the data is unchanged). The best results come from using all genes without an unsupervised embedding.

Training a linear model on top of representations derived from a linear transformation like PCA is equivalent to a regularized linear model trained on the unembedded data. Deep neural network-based architectures like SDAEs and VAEs, by contrast, process an input expression vector through a series of nonlinear transformations to learn more complex features. Therefore, we also trained a 512-dimensional SDAE and VAE on the training set for each gene set-normalization combination and used the representations derived from these neural networks as features for downstream prediction tasks. Nevertheless, we found that preprocessing the expression data using these networks decreased the out-of-sample performance of downstream prediction tasks relative to just using the normalized expression data directly (Figure 5).

### Semi-supervised representation learning

There are a variety of reasons that unsupervised representation learning can fail to discover features that are useful for downstream predictive tasks. For example, a small but consistent difference in the expression of a gene between two groups (e.g., healthy and diseased) can be used to train a highly accurate predictor. However, if this difference is much smaller than the variance in the expression of other non-predictive genes, then it will be ignored by most unsupervised representation learning algorithms. One way to avoid this problem is use a semi-supervised method to learn the representation.

The goal of semi-supervised representation learning is to derive a common set of features that are useful for multiple downstream predictive tasks. Our semi-supervised model consists of an autoencoder along with a number of logistic regression classifiers, one for each supervised task involved in the training set. The predictors operate on the 512-dimensional latent space embedding of the autoencoder. For any expression vector the autoencoder contributes a reconstruction loss. Furthermore, if there is a predictive task label associated to the expression vector, then the associated linear predictor contributes a classification loss as well. We trained the model to minimize a loss function that was a weighted combination of the autoencoder loss and the supervised loss averaged across each of the predictive tasks. We considered the out-of-sample predictive performance of four representations: the unembedded data, data embedded by a model trained using only autoencoder loss, data embedded by a model trained on the combined autoencoder and supervised losses, and data embedded by a model trained using only the supervised loss. More details are provided in the online Methods.

In order to test the semi-supervised model, we divided the larger labeled training datasets into two halves. The first half of the labeled training datasets were combined with the unlabeled data from the training set and used to train the semi-supervised autoencoder. The second half of the training datasets were held out as validation. We also held out the validation and testing labeled datasets as in the analyses of the representations learned by unsupervised algorithms. This strategy provided two types of validation tasks: those in which the representation was trained on similar data (e.g., from the same study), and those in which the representation had not been trained on similar data. The results are shown in Figure 6. Using the learned features slightly improved median predictive performance on the divided tasks but did not improve predictive performance on the validation and testing tasks used in the previous analyses.

**FIG. 6:**
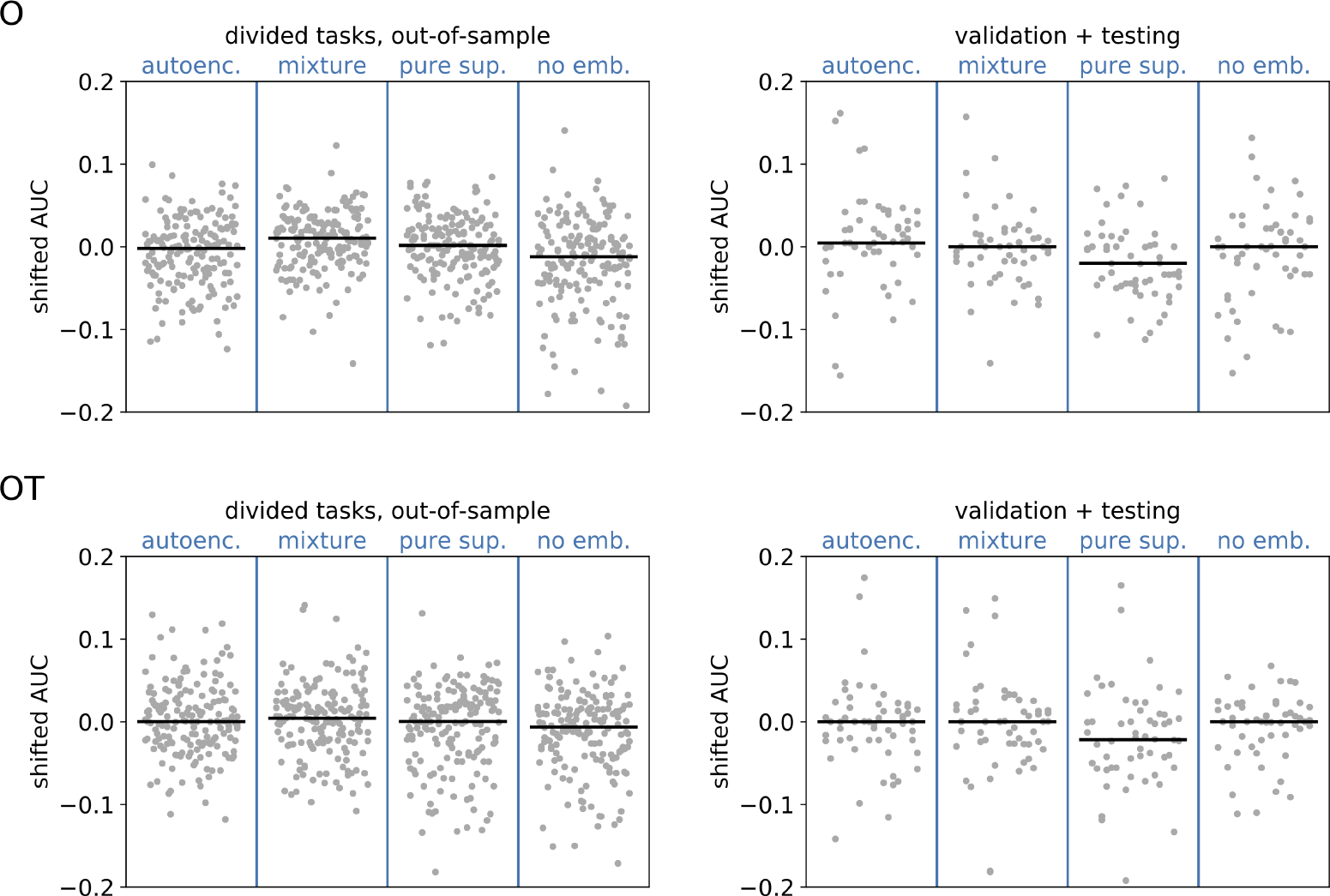
Performance for semi-supervised models. The binary task performance is shown for four different types of embedding models across two different datasets and two different gene sets. The four models are a purely unsupervised autoencoder (autoenc.), a semi-supervised embedding model (mixture), a purely supervised embedding model (pure sup.), and a no-embedding model (no emb.). To train the supervised component of the embedding models, specific task datasets are divided into two halves, one contributing to the supervised loss in training, and the other held out; the performance of the four models on the held-out halves are shown in the left column. The performance of the same models on the validation and testing tasks (which take no part in training any embedding model) are shown in the right column. Models on the **O** gene set are shown in the upper row, and on the **OT** gene set in the lower row. The gray points show the shifted AUCs on all tasks in each group and all model types, which include all supervised model types and all transforms. The bars show the median score.

## III. DISCUSSION

The hypothesis that gene expression measurements can be combined into higher level features that should be useful for predicting phenotypic characteristics has intuitive appeal. Indeed, we believe that genes act together as coordinated pathways that control cellular processes. Moreover, changes in expression at the tissue level could reflect higher level changes due to differences in cellular composition. As a result, one would expect that it is possible to define useful highlevel features for expression data; this intuition has driven the development of pathway analyses [32–34], gene set analyses [11],[10], knowledge graphs [15],[14], and cell-type deconvolution approaches [35–37] to analyzing transcriptomics experiments. More recently, a number of studies have introduced deep learning methods that aim to discover useful gene, or transcript, combinations that reflect the underlying biology without imposing particular prior knowledge [4],[5],[38– 40]. In theory, these learned representations should provide better predictive performance because they are transferring biological knowledge derived from one dataset to another. In addition, they reduce the dimension of the input data and, as a result, potentially mitigate overfitting. Here, we set out to systematically and rigorously assess the impact of these representations on downstream predictive tasks.

Our key results can be summarized in a few bullet points:

- Multivariate predictors outperformed predictors based on the best single gene.
- Larger gene sets performed better than smaller gene sets.
- CLR and tissue-specific Z-score normalization were better than TPM.
- Logistic regression and random forests performed equally well.

Representations derived from unsupervised or semi-supervised methods did not improve out-of-sample performance for phenotype prediction. Based on these key results, we conclude that *l*_2_-regularized regression applied to the CLR transformed relative transcript abundances is generally the best choice for predictive analyses using transcriptomics data. The Z-score and Z-ternary normalizations generally perform comparably to CLR, but require the GTEx data as a reference and hence CLR is recommended.

Figures 2-5 present results for the evaluation of unsupervised models on supervised tasks, studying performance as different aspects of the models change. Figure 2 shows how the performance varies across supervised tasks and demonstrates that the recommended model is nearly always one of the better performing models. Figure 3 presents the relative performance for the choices of normalization and gene set, showing that using larger gene sets improves performance on survival tasks. Figure 4 presents the performance across supervised tasks for different choices of the supervised model, showing that random forest and logistic regression models perform well. Figure 5 shows the relative performance across different unsupervised models, divided by gene set, demonstrating that supervised models on unembedded data for all genes are the best performing. All supervised evaluation results are recorded and further visualized in the appendices (Results Table and Figure 7). Taken together, these motivate the choice of the recommended model.

**FIG. 7:**
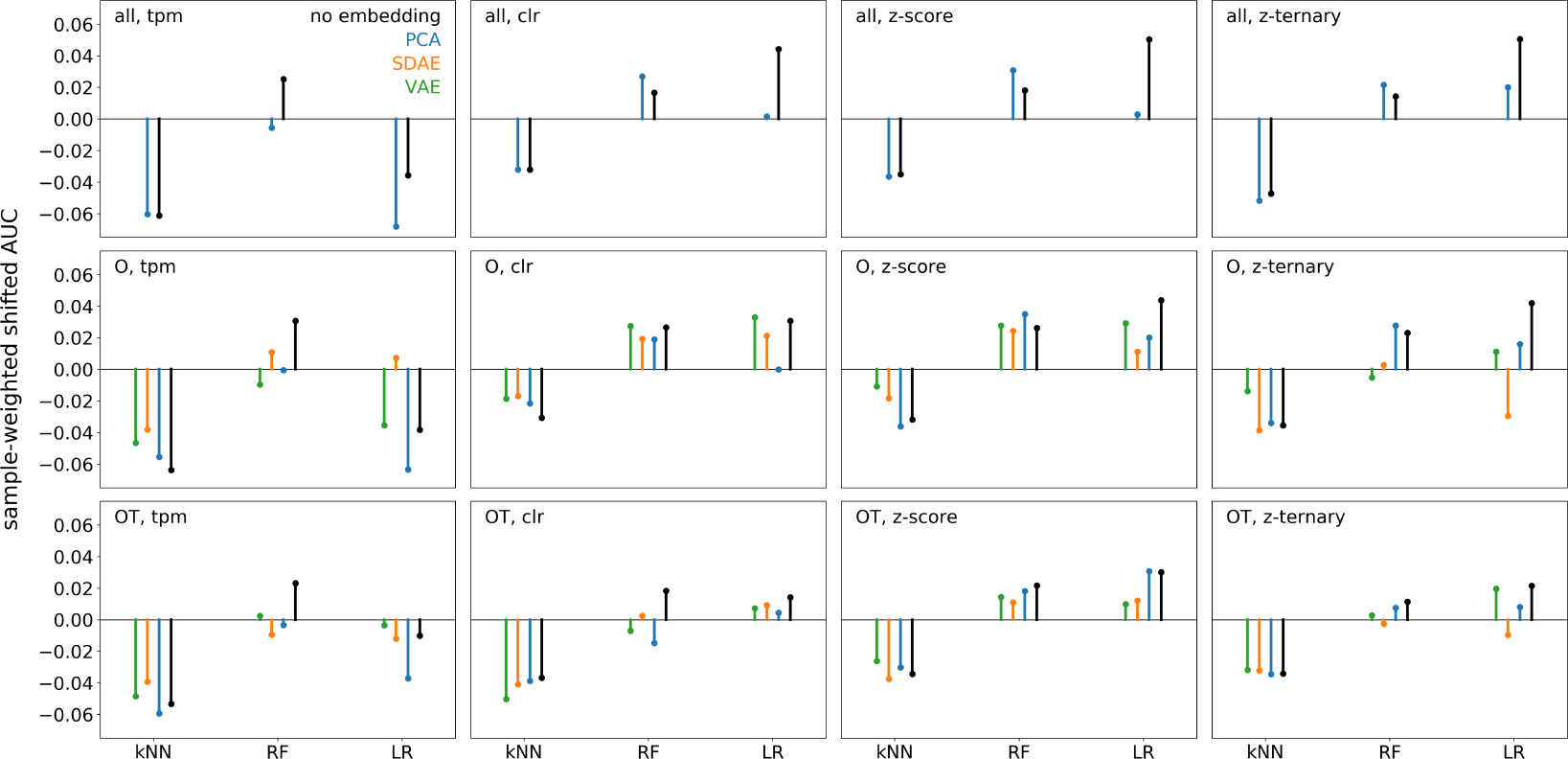
Performance over all model types. The binary task performance of each unique model type is shown for each combination of gene set, transform, supervised model, and unsupervised model. Each plot is a specific gene set and transform combination, and inside each plot results are grouped by supervised model and colored by unsupervised model. The performance shown is the average of shifted AUCs across binary tasks, weighted by the number of samples in each task to reduce the effect of fluctuations in tasks with fewer samples. The best results come from using all genes without an unsupervised embedding.

Our first conclusion, that multivariate predictors outperform predictors based on single gene expression measurements, was the clearest cut. This has some practical consequences when combined with our other conclusion that larger gene sets are better, especially for the fitting of proportional hazards models used for survival analyses. First, using multivariate predictors on large gene sets means that the number of covariates will almost always vastly outnumber the subjects in a study. Therefore, it is absolutely necessary to regularize these models by adding penalties to the coefficients. Moreover, nested cross validation should be used for all performance assessments to mitigate overfitting to hyperparameter choices and to minimize variance in the performance metric. Second, it is often impractical —or even impossible— to fit these models using standard methods on typical computing architectures. For example, open source packages for survival analyses typically use second-order methods to optimize the objective function. This works for a single gene, but fitting the multivariate model requires computing a very large matrix of second derivatives, e.g., 56,000 × 56,000 in this study. As a result, it was necessary to implement first-order optimization methods and perform most of the matrix operations using graphical processing units to make the survival analyses in this study feasible.

Overall, we found that choices such as the normalization method, the gene set, the type of supervised prediction algorithm, and the use of a learned representation made surprisingly little impact on out-of-sample predictive performance. Moreover, we could not identify any clear trends. For example, it is not necessarily better to use smaller gene sets or other lower dimensional representations for studies with smaller sample sizes. In light of these results, it is not clear that features derived from either prior knowledge or from representation learning methods have much value in the analyses of bulk RNA-seq data. If the relationship between bulk gene expression and phenotype is not one-to-one, then there is already a limit on how well one could predict phenotype from gene expression. Relatively simple methods may be already very close to this limit. Improving ‘omics-based phenotype prediction is likely to be contingent on other factors including reduction of systematic errors in sequencing data, incorporation of other data types such as single-cell sequencing and proteomics, and improved use of prior knowledge.

## IV. METHODS

The analysis presented here and depicted in Figure 1 is a multi-step procedure, starting from read counts data in the recount2 database and ending at performance metrics for various models. There are principally three stages: dataset preparation, unsupervised model training, and supervised model training.

### Dataset preparation

The recount2 database [7] is a repository of transcriptomics data sourced from over 2000 independent transcriptomics experiments. The transcriptomics data from these experiments has been reprocessed using a uniform processing pipeline, forming a single dataset amenable to large scale computational analyses. Such analyses would otherwise be problematic due to systematic differences between the original processing pipelines. The data in recount2 consists of counts of gene reads as well as exon-level quantifications. Our study concerned the gene counts data exclusively.

The data comprising recount2 can be divided into three broad groups according to their sources: GTEx, TCGA, and SRA. The GTEx group was sourced from the Genotype Tissue Expression program and contains 9538 samples from healthy individuals across 30 tissue types. The TCGA group was sourced from the Cancer Genome Atlas project and contains 11284 samples from individuals with cancer across 21 tissue types. In that group samples were taken from tumor sites as well as normal tissue adjacent to tumor (NAT) sites. GTEx and TCGA are each single, large collaboration projects with high quality control standards and protocols for sample processing. Metadata for these projects is extensive. The SRA group contains 49638 samples from 2033 smaller, distinct experiments collected in the Sequence Read Archive. Metadata for experiments in SRA are sparser, with tissue labels occasionally absent.

In total, 70460 samples were available in recount2 when the database was downloaded. However, many of these samples are not ideal for representation learning with transcriptomics data. We developed a quality control (QC) pipeline to remove samples or entire SRA studies. The number of samples remaining after the QC pipeline is 39848. The QC steps are as follows:

- Remove samples in which the reported cell type is a cell line. 9644 samples fit this criterion.
- Remove studies in SRA from single-cell sequencing. Examining metadata from GEO, 38 studies in SRA with 5865 samples in recount2 have single-cell transcriptomic data.
- Remove samples in which the reported tissue does not match any tissue in GTEx. 6824 samples fit this criterion (see Z-score normalization later).
- Remove samples in SRA which have duplicate GEO accession numbers (GSMs). There were 9601 samples that met this criterion.
- Remove samples in which more than 30% of genes listed in the Gene Ontology (GO) 12 under the “biological process” or “molecular function” categories have a counts value of 0. 15390 samples met this criterion.

The number of matching samples in each step are non-exclusive, meaning a sample can match more than one of the exclusion criteria. These effect of these exclusion criteria are depicted in the supplementary figures (Figure 8). In total we removed 30612 samples, approximately 43% of the total. No GTEx samples were removed; and only 521 TCGA samples were removed.

**FIG. 8:**
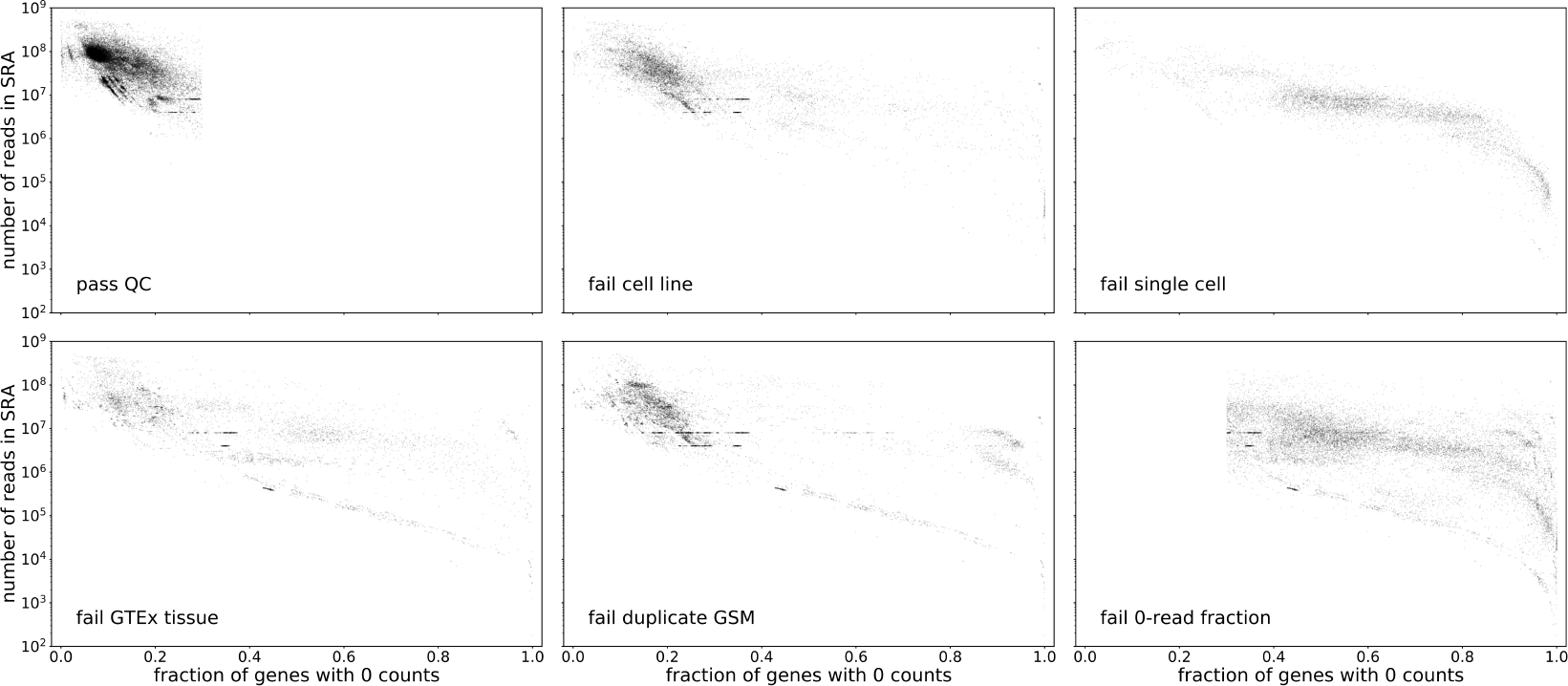
Effect of the quality control cuts. Gene expression samples plotted in terms of two quality metrics, the number of reads and the fraction of genes with zero reads. The upper left plot shows each sample in the recount2 database passing quality cuts. The remaining plots show samples failing each of the quality cuts. The cuts remove a swath of samples whose characteristics are distinct from the bulk of high-quality samples retained in the dataset.

**FIG. 9:**
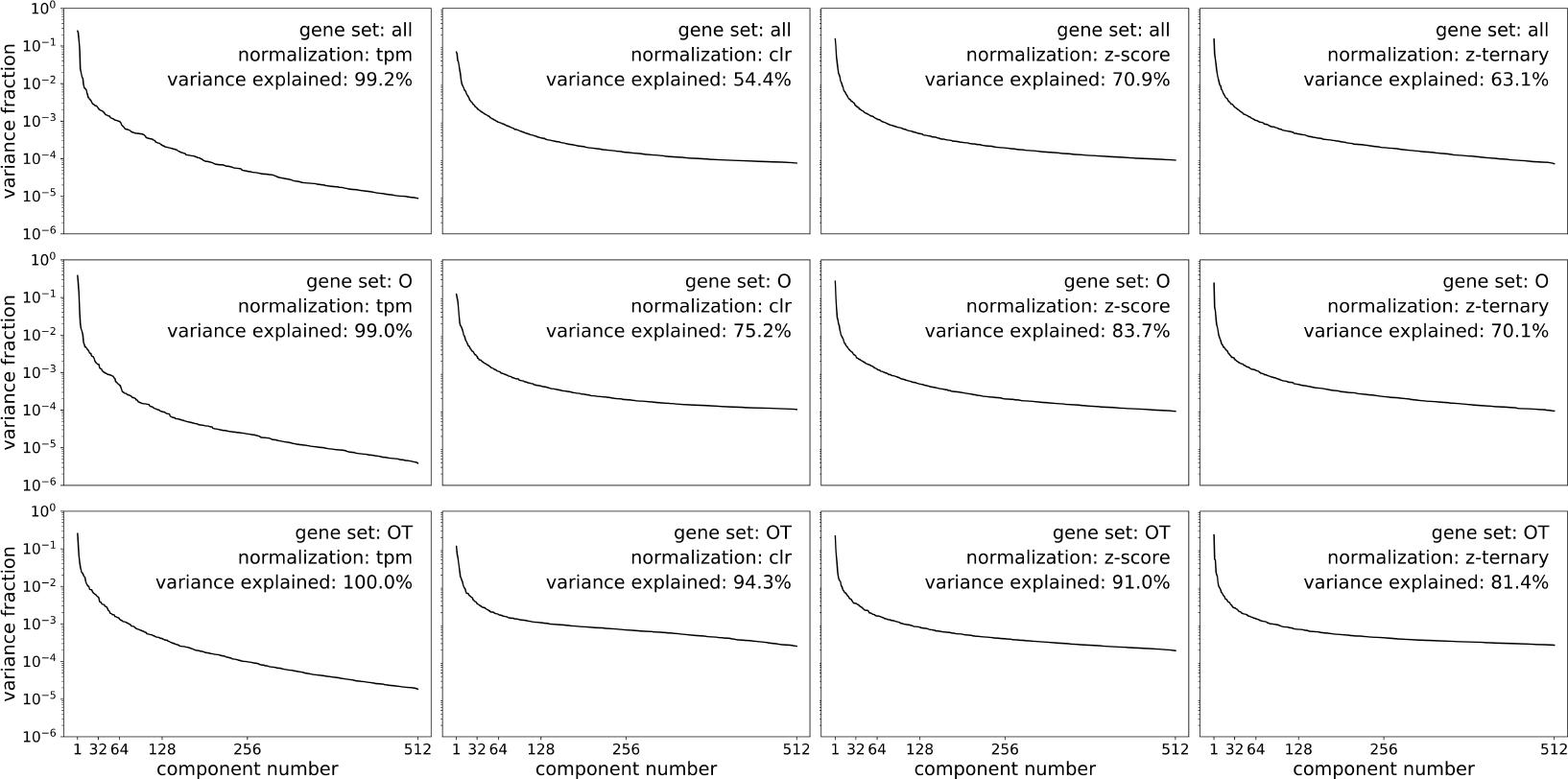
Variance explained by PCA. Per-component variance explained by PCA for each gene set and normalization combination. The total variance explained by all 512 components is displayed on each plot. The majority of variance is captured by the PCA in all cases; in some nearly all variance is captured.

It is useful to present some detailed commentary on the duplicate GEO accession number criterion. We observed that several samples have duplicate GSMs, and that many such samples had the same number of reads (a round number, e.g., 8 million). This suggests that the individual samples could be chunks of reads from the same underlying sample. However, we could find no satisfying reason for duplicate GSMs or the round number of read counts for these samples, and therefore excluded them from the dataset.

The QC pipeline determines which samples are admitted to the final dataset; there are also choices to be made about which genes to consider in the analysis, and which normalizing procedures to apply to the expression data.

We considered three different gene sets in the analysis:

- **all** genes: (57992 genes).
- **O** genes: genes in GO under the “biological process” or “molecular function” categories (17970 genes at the time of dataset creation).
- **OT** genes: **O** genes also labeled as transcription factors 29 (1530 genes at the time of dataset creation).

In addition, we considered four different normalizing transformations of the counts data:

- **TPM**: The counts are transformed into transcripts per million (tpm), which account for gene length to normalize reads. The TPM value is determined in terms of the counts as,

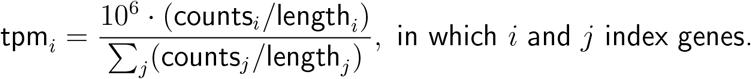
- **CLR**: A centered-log-ratio transform is carried out on the TPM vectors. The CLR value is determined in terms of the TPM values as,

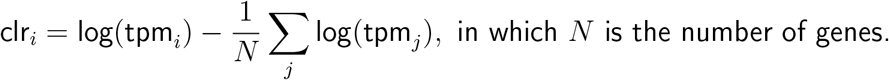
- **Z-score**: The Z-score transform is carried out on the CLR features. The Z-score is the CLR value standardized by the mean expression of a gene in healthy tissue, determined by the GTEx samples for the same tissue. The Z-score value is determined in terms of the CLR value as,

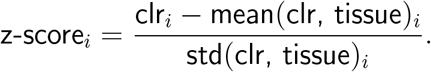
- **Z-ternary**: The Z-ternary transform is carried out on the Z-score features. The Z-score values are ternarized based on their value, and the ternarization indicates whether the gene’s expression is increased, decreased, or unchanged relative to the mean expression in healthy tissue. Since the distribution of Z-score values is expected to be approximately normal for healthy tissue, any Z-score value below −2 is assigned the Z-ternary value of −1; any Z-score value above 2 is assigned the Z-ternary value of 1, and any Z-score value between −2 and 2 is assigned the Z-ternary value of 0.

We made use of the open-source python library *genemunge* [41] for making these normalizations and selecting the gene sets. Each of the normalizations are carried out on the expression data for all genes. Whenever a smaller gene set is used, the values of the features for the selected genes are simply taken from the data for all genes. The three gene sets and four normalizations yield twelve different datasets that are used in the analysis.

### Tasks and dataset allocation

The above procedure describes the preparation of the gene expression datasets. In addition to the expression data, some samples have one or more labels suitable for predictive modeling. TCGA has rich metadata with natural label types, available in the TCGA Pan-Cancer Clinical Data Resource [23]; some SRA studies also contain useful metadata in GEO [22] relevant to human disease. From the TCGA and recount2 metadata we selected four categories of predictive tasks: binary labels for the grade of a tumor in various cancer types (8 tasks); binary labels for the stage of a tumor in various cancer types (10 tasks); times for overall survival in various cancer types (13 tasks); and times for progression free interval in various cancer types (13 tasks). From the GEO metadata we selected binary and multiclass labels for various clinical characteristics (6 tasks) [42–47]. In total there are 50 tasks for which supervised models may be built.

We divided the 50 predictive tasks into three groups, “training”, “validation”, and “test”. We then built a “training” gene expression dataset consisting of any samples with a label in the training task group, as well as any samples with no label. This dataset, which has 36794 gene expression samples, was used to train the unsupervised models. A sample’s inclusion in this dataset distinguishes the training and validation task groups. Both the training and validation tasks were used in the analysis, whereas tasks in the test group were held out until the end of the project so that no model selection criteria might influence performance on these tasks in any way. The supervised tasks are summarized in the appendix (Tables I,II,III,IV).

**Table I:** TCGA binary tasks. The 18 binary tasks derived from TCGA used to train supervised models and validate the unsupervised embeddings. The tasks are grouped into two categories, TCGA tumor stage tasks (10), and TCGA tumor grade tasks (8). The project names correspond to those in Figure 2.

| **TCGA stage tasks** |  |  |  |  |  |
| --- | --- | --- | --- | --- | --- |
| project | disease | label | label type | group | samples |
| COAD | colon adenocarcinoma | II- vs. III+ | binary | train | 505 |
| KIRC | clear cell renal cell carcinoma | II- vs. III+ | binary | train | 544 |
| LIHC | hepatocellular carcinoma | I- vs. II+ | binary | train | 374 |
| LUAD | lung adenocarcinoma | I- vs. II+ | binary | train | 542 |
| SKCM | cutaneous melanoma | II- vs. III+ | binary | train | 249 |
| STAD | stomach adenocarcinoma | II- vs. III+ | binary | train | 416 |
| THCA | thyroid cancer | I- vs. II+ | binary | train | 513 |
| UCEC | uterine corpus endometrial carcinoma | I- vs. II+ | binary | train | 554 |
| LUSC | lung squamous cell carcinoma | I- vs. II+ | binary | validate | 504 |
| BRCA | breast invasive carcinoma | II- vs. III+ | binary | test | 1134 |
| **TCGA grade tasks** |  |  |  |  |  |
| project | disease | label | label type | group | samples |
| CESC | cervical squamous cell carcinoma | II- vs. III+ | binary | train | 306 |
| KIRC | clear cell renal cell carcinoma | II- vs. III+ | binary | train | 544 |
| LGG | low grade glioma | II- vs. III+ | binary | train | 532 |
| LIHC | hepatocellular carcinoma | II- vs. III+ | binary | train | 374 |
| PAAD | pancreatic adenocarcinoma | II- vs. III+ | binary | train | 179 |
| STAD | stomach adenocarcinoma | II- vs. III+ | binary | train | 416 |
| UCEC | uterine corpus endometrial carcinoma | II- vs. III+ | binary | train | 554 |
| HNSC | head-neck squamous cell carcinoma | II- vs. III+ | binary | test | 504 |

**Table II:** SRA tasks. The 8 tasks derived from SRA used to train supervised models and validate the unsupervised embeddings. The project names correspond to those in Figure 2.

| **SRA tasks** |  |  |  |  |  |
| --- | --- | --- | --- | --- | --- |
| project | disease | label | label type | group | samples |
| GSE65832 | atopic dermatitis | lesional vs. not | binary | train | 40 |
| GSE66207 | Crohn’s disease | type: B1, B2 or B3 | multiclass (3) | train | 20 |
| GSE72819 | ulcerative colitis | treatment remission | binary | validate | 69 |
| GSE47944 | psoriasis | lesional vs. not | multiclass (3) | validate | 63 |
| GSE50244 | diabetes | normoglycemic, impaired, diabetic | multiclass (3) | validate | 76 |
| GSE67785 | psoriasis | lesional vs. not | binary | test | 28 |

**Table III:** TCGA overall survival (OS) tasks. The 13 overall survival tasks derived from TCGA used to train supervised models and validate the unsupervised embeddings. The project names correspond to those in Figure 2.

| **TCGA OS tasks** |  |  |  |  |  |
| --- | --- | --- | --- | --- | --- |
| project | disease | label | label type | group | samples |
| CESC | cervical squamous cell carcinoma | OS | survival | train | 304 |
| COAD | colon adenocarcinoma | OS | survival | train | 455 |
| ECSA | esophageal carcinoma | OS | survival | train | 184 |
| KIRP | papillary renal cell carcinoma | OS | survival | train | 289 |
| LUAD | lung adenocarcinoma | OS | survival | train | 507 |
| OV | ovarian cancer | OS | survival | train | 420 |
| PAAD | pancreatic adenocarcinoma | OS | survival | train | 178 |
| SARC | sarcoma | OS | survival | train | 259 |
| STAD | stomach adenocarcinoma | OS | survival | train | 409 |
| UCEC | uterine corpus endometrial carcinoma | OS | survival | train | 540 |
| HNSC | head-neck squamous cell carcinoma | OS | survival | validate | 501 |
| BLCA | urothelial bladder carcinoma | OS | survival | test | 407 |
| LUSC | lung squamous cell carcinoma | OS | survival | test | 495 |

**Table IV:** TCGA progression-free interval (PFI) tasks. The 13 progression-free surivival tasks derived from TCGA used to train supervised models and validate the unsupervised embeddings. The project names correspond to those in Figure 2.

| **TCGA PFI tasks** |  |  |  |  |  |
| --- | --- | --- | --- | --- | --- |
| project | disease | label | label type | group | samples |
| CESC | cervical squamous cell carcinoma | PFI | survival | train | 304 |
| COAD | colon adenocarcinoma | PFI | survival | train | 455 |
| ECSA | esophageal carcinoma | PFI | survival | train | 184 |
| KIRP | papillary renal cell carcinoma | PFI | survival | train | 288 |
| LUAD | lung adenocarcinoma | PFI | survival | train | 507 |
| OV | ovarian cancer | PFI | survival | train | 420 |
| PAAD | pancreatic adenocarcinoma | PFI | survival | train | 178 |
| SARC | sarcoma | PFI | survival | train | 259 |
| STAD | stomach adenocarcinoma | PFI | survival | train | 411 |
| UCEC | uterine corpus endometrial carcinoma | PFI | survival | train | 540 |
| HNSC | head-neck squamous cell carcinoma | PFI | survival | validate | 501 |
| BLCA | urothelial bladder carcinoma | PFI | survival | test | 408 |
| LUSC | lung squamous cell carcinoma | PFI | survival | test | 496 |

**Table V:** SDAE Architectures and Hyperparameters. - All data was standardized before training.
- Weights were intialized randomly according to a central Gaussian distribution of standard deviation `init. weight std. dev`..
- The learning rate was reduced from `init. LR` by a factor of `LR gamma` every `LR` step epochs.
- All activations were ReLU except for the final layer, which was *linear* or *hardtanh* in the case of Z-ternary normalization.
- We saw no perceived benefit from *l*_2_-regularization over-against selection of the noise level, and so simply fixed the *l*_2_ coefficient *l*_2_-coeff. to 0. The value of noise std. dev. was selected by assessing validation performance over a range of values from 0 to 0.5.
- All SGD used ADAM [56] with parameters (0.5, 0.999).
- Models were first trained in a greedy-layerwise fashion before being trained end-to-end. Both training eras used the same set of hyperparameters.

| gene set | layer dimensions |
| --- | --- |
| O | 17970 - 2048 - 1024 - 512 - 1024 - 2048 - 17970 |
| OT | 1530 - 1530 - 1024 - 512 - 1024 - 1530 - 1530 |

TABLE V: **SDAE Architectures and Hyperparameters.**
| init. weight<br>std. dev. | noise<br>std. dev. | epochs | batch<br>size | init. LR | LR step | LR gamma | $l_2$-coeff. |
| --- | --- | --- | --- | --- | --- | --- | --- |
| 0.01 | 0.3 | 500 | 50 | 0.0001 | 50 | 0.8 | 0 |

**Table VI:**
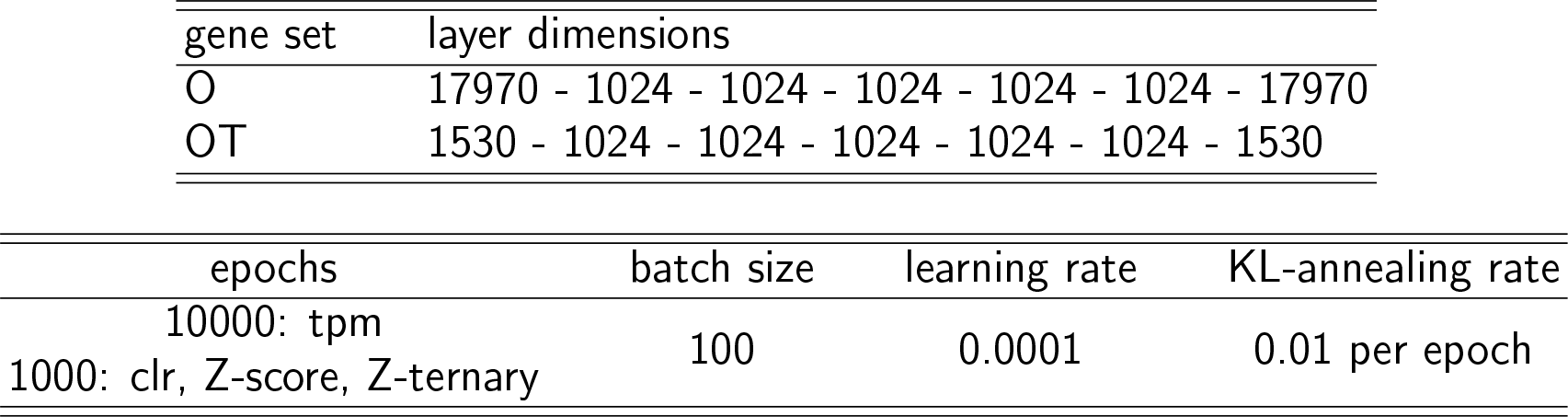
VAE Architectures and Hyperparameters. - Weights were intialized with a centered Gaussian distribution with a standard deviation equal to the inverse of the number of input features.
- Models were trained with the KL-annealing rate [52] moving from 0 to 1 linearly over the first 100 epochs.
- The learning rate was held constant in training.
- No additional regularization was performed due to a strong correlation between training and validation loss.

**Table VII:** Logistic Regression Hyperparameters. - Weights were initialized randomly according to the standard (Xavier) Glorot normal [57] prescription.
- The learning rate was reduced from `init. LR` by a factor of `LR gamma` every LR step epochs.
- The *l*_2_-coeff value is selected by cross-validation over the range 10^*−*6^ to 10^3^ in logarithmic steps of 10.
- All SGD used ADAM [56] with parameters (0.5, 0.999).

| epochs | batch size | init. LR | LR step | LR gamma |
| --- | --- | --- | --- | --- |
| 200: binary<br>300: multiclass | $\text{floor}(\text{num. samples} / 10)$ | 0.001 | 10 | 0.9 |

**Table VIII.** Cox Proportional Hazards Hyperparameters. - The neural network model consists of a batch-normalization layer [58] followed by a single, linear fully-connected to compute the relative risk function.
- Weights were initialized randomly according to the standard (Xavier) Glorot normal [57] prescription.
- The learning rate was held constant during training.
- The *l*_2_-coeff value is selected by cross-validation over the range 10^*−*6^ to 10^3^ in logarithmic steps of 10.
- All SGD used ADAM [56] with parameters (0.5, 0.9).

| epochs | batch size | init. LR |
| --- | --- | --- |
| 500 | $\text{floor}(\text{num. samples} / 5)$ | 0.00001 |

**Table IX.**
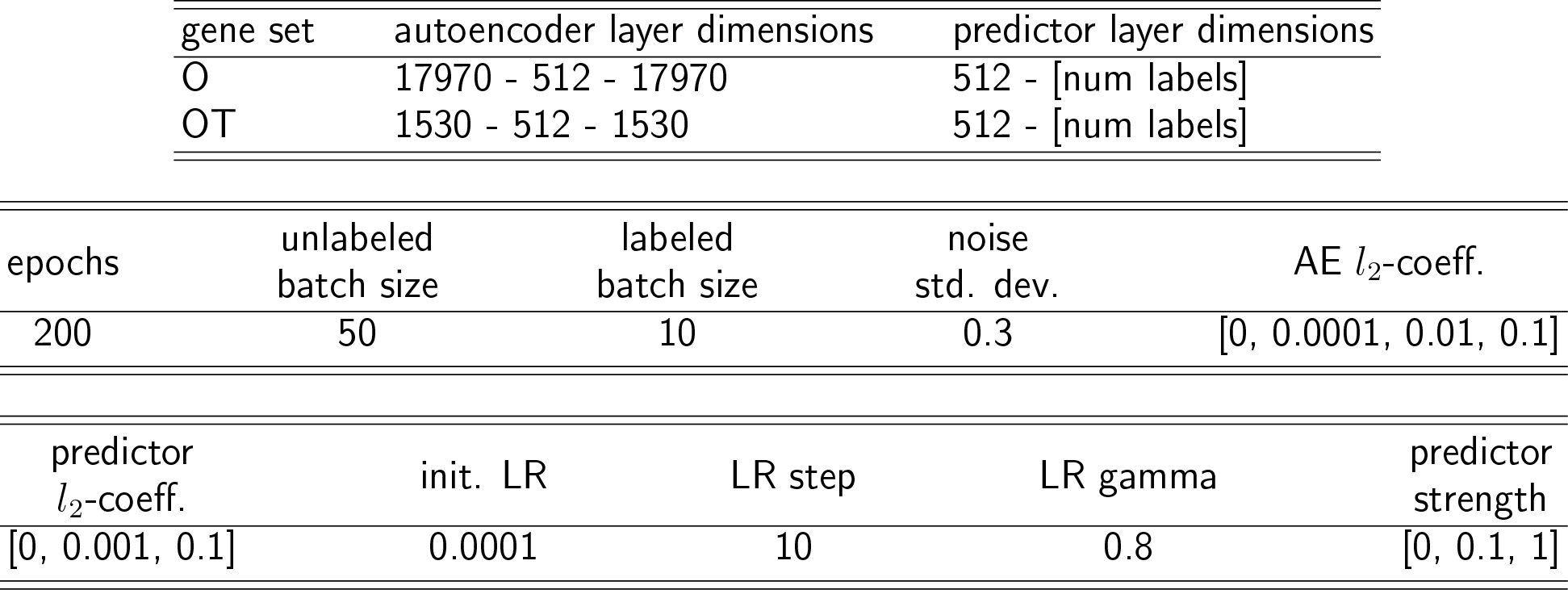
Semi-supervised Model Architectures and Hyperparameters. - All data was standardized before training.
- Brackets indicate different possible values. All possible combinations of parameters were tried with the best parameter set chosen by virtue of performance on held-out half of the divided predictive tasks.
- Weights were initialized randomly according to a central Gaussian distribution with standard deviation of 0.01.
- The learning rate was reduced from `init. LR` by a factor of `LR gamma` every `LR step` epochs.
- All SGD used ADAM [56] with parameters (0.9, 0.9).

### Unsupervised models

Three different types of unsupervised models were trained on the gene expression datasets: principal components analysis (PCA), stacked denoising autoencoders (SDAE), and variational autoencoders (VAE).

#### Principal components analysis (PCA)

The problem of finding the kprincipal components of a suitably large collection of vectors of dimension n admits an analytic solution. But the computation required to perform this calculation is in *O*(*n*^3^), making it intractable in high dimensions. Due to the high dimension of the larger gene sets (*>*17k), we performed the PCA analysis via stochastic gradient descent following the algorithm introduced by Arora et. al. [48] called “Stochastic Approximation.” The leading 512 principal components were retained.

#### Stacked denoising autoencoders (SDAE)

We employed denoising autoencoder architectures of “hourglass” shape with seven layers. The hourglass narrows to a middle layer of 512 dimensions, yielding a 512-dimensional encoder. The details of the architecture are recorded in the appendix. The models were trained with stochastic gradient descent to minimize the mean squared reconstruction loss. We found that pre-training the models layerwise before end-to-end training produced the best results. Therefore these models are best described as stacked denoising autoencoders per the original presentation [49]. The models were regularized by input noise variance and an *l*_2_ weight penalty, with these hyperparameters selected by sweeping a range.

#### Variational autoencoders (VAE)

We also included a deep generative model among our unsupervised model types, the variational autoencoder [50]. In particular, we employed the methods of Klambauer, et. al. [51] which make use of self-normalizing units, SNNs, for improved training dynamics and representational capability. We trained the models using the KL-annealing method of Bowman et. al. [52] during the first 100 epochs and then let training proceed with the normal loss function for the remaining epochs. The layer dimensions are recorded in the appendix. The latent encodings consist of 512 dimensions for the distributional means and 512 for the distributional log variances. Therefore the trained model’s feature encoder is the restriction to the 512 dimensions of the means variables.

#### The “no-embedding” model

In addition to these unsupervised models, we also employed a kind of control comparator: a “no-embedding” model which does nothing to the expression data. The dimension of the gene expression data is not reduced under the no-embedding model; the features are the normalized gene expression vectors themselves.

#### Computational constraints

We trained the PCA model on each of the four normalizations for each of the three gene sets. Due to computational constraints we applied the SDAE and VAE models to each of the four normalizations for the **O** and **OT** gene sets excluding the all genes set. The lack of an improvement in performance on smaller gene sets indicated the dataset with all genes was unlikely to provide quality embedding models.

### Supervised models

We evaluated the ability of a unsupervised model to learn useful representations across transcriptomics data by assessing the performance of supervised models operating on the learned representations. For each unsupervised model and predictive task, we trained and evaluated supervised models using nested cross validation. The performance of these predictive models gave an indication of how well the learned representation captured features in the data useful for various kinds of phenotype prediction. Before presenting the different kinds of supervised models, we present a small primer on nested cross validation.

### Nested cross validation

Nested cross validation is designed to provide a robust estimate of the expected (predictive) model performance on new data, optimizing over a set of hyperparameter values (such as the maximum depth in a random forest). In nested cross validation, there are two loops over the data, the outer and inner loop. The inner loop is used to select an optimal hyperparameter value, and the outer loop is used to estimate the performance of the model with this hyperparameter value. In the outer loop, data is divided evenly into *K* groups, or folds (we use *K* = 5). For each fold, the data for that fold is held out and the remaining *K −* 1 folds are used for the inner loop. In the inner loop, this data is divided into *K* folds, and on each fold the data for that fold is held out and the model is trained on the remaining *K −* 1 folds for each hyperparameter value. The held-out fold is used to estimate the model performance for each hyperparameter value, and this performance is averaged over all folds in the inner loop. The best performing hyperparameter value is selected, and the model is re-trained on all data used in the inner loop. The model performance is then evaluated on the held out data from the outer fold. This value is averaged over all folds in the outer loop, and this final average is the estimated model performance. Note that a different optimal hyperparameter may be selected for each outer fold. Nested cross validation is resistant to hyperparameter overfitting, as the model is evaluated on data completely held out from the process of selecting the optimal hyperparameter. With this robustness comes increased computational complexity —if there are *N* hyperparameter values tested, nested cross validation requires training *K*(*KN* + 1) individual models.

#### Classification tasks

For classification tasks we applied three different types of supervised models:

- **Logistic regression (LR)**. Logistic regression with an l2 penalty, trained via stochastic gradient descent. The logistic regression model is a single layer neural network with a softmax activation on the output. The hyperparameter optimized was the *l*_2_ penalty, logarithmically spaced between 10^*−*6^ and 10^3^ in ten steps. The model was implemented in pytorch [53]. Hyperparameters and training notes are provided in the appendix.
- **Random forest (RF)**. Random forest models with 100 trees per forest. The hyperparameter optimized was the maximum depth of the random forest, logarithmically spaced between 2 and 2^7^ in seven steps. We relied on the scikit-learn [54] implementation of random forest.
- **K-nearest neighbors (kNN)**. The hyperparameter optimized was the value of k, the number of neighbors used, taking a value of 1, 3, 5, 7, or 9.

#### Survival tasks

For survival tasks, in which the overall survival time or the progression free interval time were predicted, we trained a Cox proportional hazard (CPH) model. The standard solvers for CPH models use second-order methods, such as versions of Newton’s method, making them unsuitable for use with a large number of features. The computation time required for the 512-dimensional embedding, using nested cross validation, is already immense; training CPH models on data without an embedding is completely impractical. Instead, we implemented a CPH model in pytorch, and trained it via stochastic gradient descent by backpropagating through the Cox-Efron pseudolikelihood[55]. Such models can be trained with a large number of features —even all genes— and can be GPU accelerated. We regularized these models with an *l*_2_ penalty whose strength, logarithmically spaced between 10^*−*6^ and 10^3^ in ten steps, was optimized in the inner cross-validation loop. Even with this computational speedup, evaluating the survival tasks requires the bulk of compute time. It bears noting that these models were still trained with a fixed initial learning rate which was small enough to guarantee controlled gradient descent across all tasks. It is certain that absolute performance on individual contrasts could be improved by also optimizing the learning rate in the nested cross validation. However, because the study concerns the relative performance of this algorithm across gene sets, embeddings, and normalizations, we avoided this additional multiplier on the computational time.

#### Single-gene comparators

All of the above supervised models are trained on features from multiple genes. In order to compare our embedding models to single-gene analysis, we also trained a set of models on single genes with no-embedding model. For these models the hyperparameter optimized in the inner cross validation loop is the gene selected for the model. We had no need to run these comparators across all transform/predictor combinations so we restricted these examples to clr-transformed data and used only the univariate logistic regression models for classification tasks. For survival tasks, single-gene CPH models were trained. No regularization term was necessary in either case because these models have so few parameters. These results provide a direct comparison to the multi-gene, CLR-transformed, no-embedding results.

#### Components of supervised model results

In total, we evaluated a very large number of (multi-gene) supervised models. There are five different characteristics of a single result:

- **Task**. There are 24 classification tasks and 26 survival tasks.
- **Gene set**. There are three gene sets, **all** genes, **O** genes, and **OT** genes.
- **Normalization**. There are four data normalizations, TPM, CLR, Z-score, and Z-ternary.
- **Unsupervised model**. There are four types of unsupervised model, PCA, SDAE, VAE, and no-embedding. SDAE and VAE were only trained on the **O** and **OT** gene sets.
- **Supervised model**. For classification tasks, three different supervised models were trained, LR, RF, and kNN. For survival tasks, a CPH model was trained.

This amounts to 3920 results. In terms of individual models trained during nested cross validation, there are 807600 models.

### Semi-supervised models

The semi-supervised models are designed to learn a feature embedding which is co-adapted to the purpose of reconstruction as well as the performance of supervised models operating on the embedding. Each of the semi-supervised models consists of a linear, single-hidden layer autoencoder coupled to a number of logistic regression predictors —one for each supervised task involved in the training dataset. The supervised predictors operate on the autoencoder’s 512-dimensional encoding. Both a schematic diagram of the model architecture and the details of the architectures are recorded in the supplementary figure (Figure 10).

**FIG. 10:**
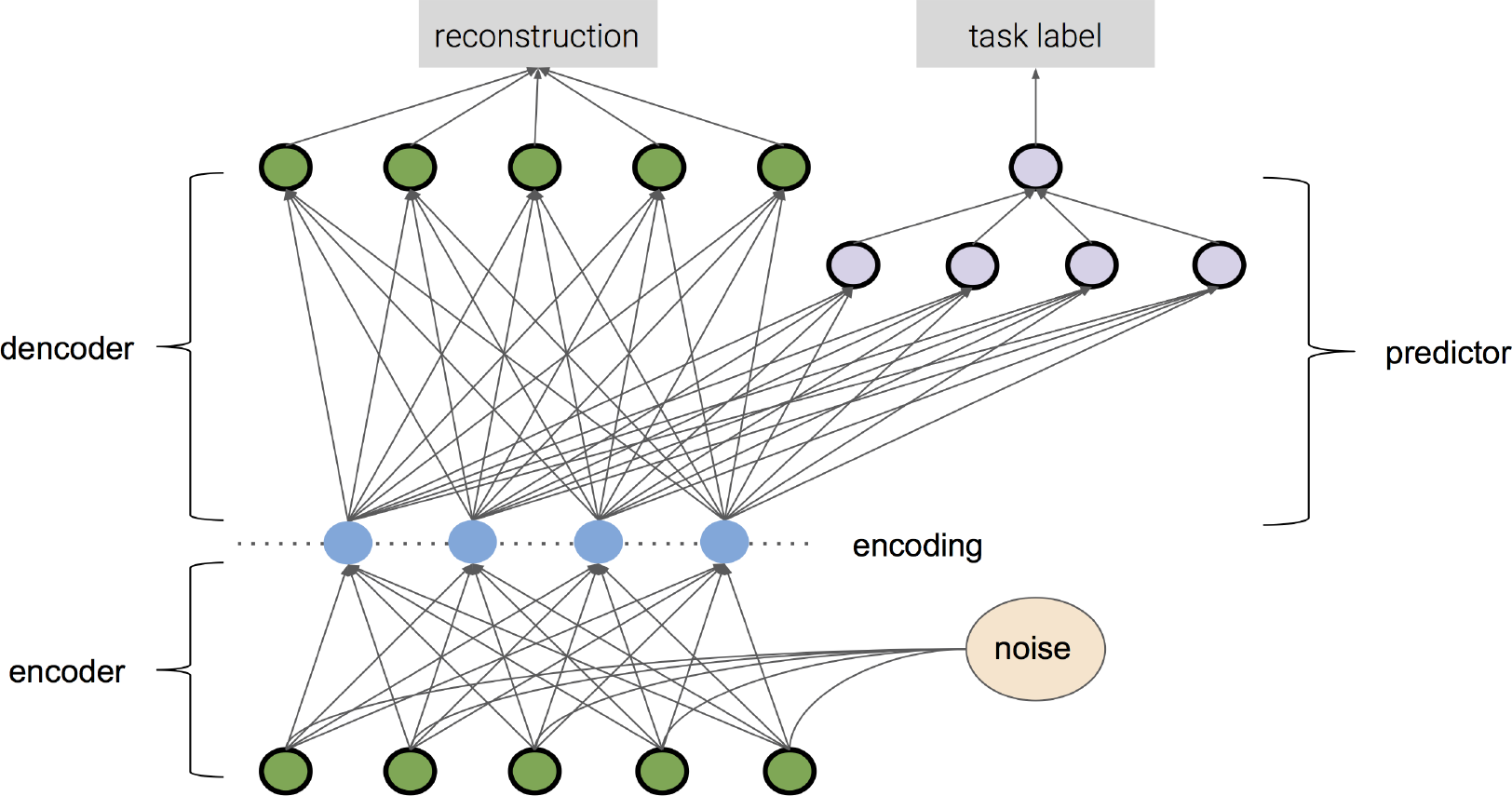
Semi-supervised model schematic. The semi-supervised model consists of a denoising autoencoder coupled to one or more predictors. The training loss is a combination of reconstruction error and classification error.

#### Data preparation

In order to be able to assess the performance of semi-supervised representation learning within-task, we had to further subdivide some of the labeled expression data. In particular, we subdivided into two halves the binary predictive tasks within the “training set” which contained at least 200 samples. The first half was used in the training of the semi-supervised model; the second half was held out for validation. We called these sets the “divided tasks”; they were drawn from the following binary tasks:

{CESC grade, COAD stage, KIRC grade, KIRC stage, LGG grade, LIHC stage, LIHC grade, LUAD stage, SKCM stage, STAD stage, STAD grade, THCA stage, UCEC stage, UCEC grade}.

The rest of the tasks which constituted the original “validation” and “test” sets were used for validation. So the training set for each semi-supervised model consisted of all expression data from the first halves of the fourteen divided tasks along with their associated binary labels.

#### Training of semi-supervised models

Given any sample expression vector *x* from the training set we can compute the autoencoder reconstruction loss on that sample, specifically as the squared reconstruction error,

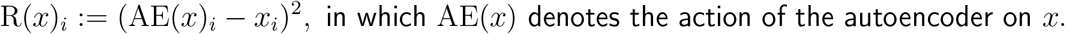

Supposing that *x* has a class label *l*_*x*_ ∈ {0, 1} from the *j*^th^ predictive task, we can also compute a classification error of the associated binary logistic regression classifier P_*j*_,

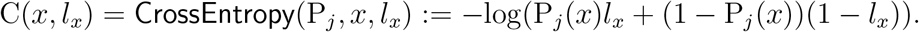

We trained our semi-supervised models (via stochastic gradient descent) to minimize a convex combination of these two error terms. The constant controlling the interpolation of these two losses we called the “predictor strength,” *π*, which ranged from 0 to 1. Our training algorithm allowed different batch sizes for the autoencoder loss and the predictor losses; let these be denoted by BR, and BC, respectively. Let *{x_m_}, {x_n_, l_xn_}* be batches drawn randomly from the training data, the first consisting of only expression vectors, the second containing both expression vectors and paired class labels. Our loss term takes the form,

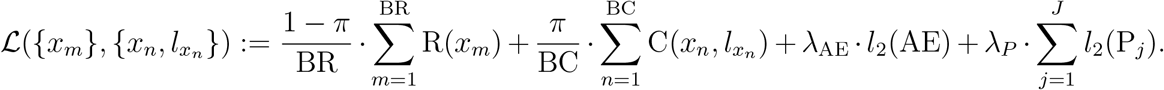

Here, *J* denotes the total number of predictive tasks. The last two terms are *l*_2_ weight penalties on the model parameters; these are controlled by adjustable constants *λ*_AE_ and *λ*_P_.

We compared three scenarios for the predictor strength in our analysis,

- *π* = 0.0, i.e. the model is an autoencoder only.
- *π* = 0.1, the model is trained with a mixture of both losses.
- *π* = 1.0, i.e. the model is a purely supervised shared-embedding model.

We also compared results to a no-embedding model as a kind of control.

For each gene set, data normalization, and predictor strength scenario, we performed a sweep over all 16 pairs of values for *λ*_AE_ and *λ*_P_ in the cartesian product *{*0, 0.1, 0.01, 0.001*}*s^2^. We selected the *l*_2_ coefficient pair which minimized average error on the held-out half of the divided contrasts.

Finally, we assessed the performance of predictive models (across all three types, LR, RF, kNN) operating on the learned data embedding to compare the effect of semi-supervised representation learning across these three scenarios. Those results are displayed in Figure 6 in the main text.

## DATA AVAILABILITY

The gene expression data used to train the unsupervised models and the gene expression data and labels used to train and evaluate the supervised models are available on figshare, and can be accessed at datasets. These data include the gene expression data for all three gene sets and all four normalizations, and the accompanying supervised labels.

## CODE AVAILABILITY

The code used to process the data and train the unsupervised, supervised, and semi-supervised models is a combination of open source and proprietary, licensable software. The open source software made available on GitHub, accessible at representation learning for transcriptomics, is a framework to carry out nested cross validation and includes an example to train the recommended model on a particular supervised task.

## AUTHOR CONTRIBUTIONS

D.Z. and C.K.F. conceived of the analysis. A.M.S., J.R.W., and J.L. carried out the analysis. A.M.S., J.R.W., and C.K.F. wrote the manuscript. All authors participated in the design of the analysis, the interpretation of its results, and revisions to the manuscript.

## References

[1] N. S. Madhukar and O. Elemento, Bioinformatics Approaches to Predict Drug Responses from Genomic Sequencing, Methods in Molecular Biology (Clifton, N.J.) 1711, 277 (2018).

[2] S. Li, P. P. Labaj, P. Zumbo, P. Sykacek, W. Shi, L. Shi, J. Phan, P.-Y. Wu, M. Wang, C. Wang, D. Thierry-Mieg, J. Thierry-Mieg, D. P. Kreil, and C. E. Mason, Detecting and correcting systematic variation in large-scale RNA sequencing data, Nature Biotechnology 32, 888 (2014).

[3] P. A. C. ’t Hoen et al., Reproducibility of high-throughput mRNA and small RNA sequencing across laboratories, Nature Biotechnology 31, 1015 (2013).

[4] Ching et al., Opportunities and obstacles for deep learning in biology and medicine, Journal of The Royal Society Interface 15, 20170387 (2018).

[5] P. Mamoshina, A. Vieira, E. Putin, and A. Zhavoronkov, Applications of Deep Learning in Biomedicine, Molecular Pharmaceutics 13, 1445 (2016).

[6] A. C. Frazee, B. Langmead, and J. T. Leek, ReCount: A multi-experiment resource of analysis-ready RNA-seq gene count datasets, BMC Bioinformatics 12, 449 (2011).

[7] L. Collado-Torres, A. Nellore, K. Kammers, S. E. Ellis, M. A. Taub, K. D. Hansen, A. E. Jaffe, B. Langmead, and J. T. Leek, Reproducible RNA-seq analysis using recount2, Nature Biotechnol-ogy 35, 319 (2017).

[8] A. Lachmann, D. Torre, A. B. Keenan, K. M. Jagodnik, H. J. Lee, L. Wang, M. C. Silverstein, and A. Ma’ayan, Massive mining of publicly available RNA-seq data from human and mouse, Nature Communications 9, 1366 (2018).

[9] S. E. Ellis, L. Collado-Torres, A. Jaffe, and J. T. Leek, Improving the value of public RNA-seq expression data by phenotype prediction, Nucleic Acids Research 46, e54 (2018).

[10] M. Gönen, Integrating gene set analysis and nonlinear predictive modeling of disease phenotypes using a Bayesian multitask formulation, BMC Bioinformatics 17, 0 (2016).

[11] A. Subramanian, P. Tamayo, V. K. Mootha, S. Mukherjee, B. L. Ebert, M. A. Gillette, A. Paulovich, S. L. Pomeroy, T. R. Golub, E. S. Lander, and J. P. Mesirov, Gene set enrichment analysis: A knowledge-based approach for interpreting genome-wide expression profiles, Proceedings of the National Academy of Sciences 102, 15545 (2005).

[12] M. Ashburner, C. A. Ball, J. A. Blake, D. Botstein, H. Butler, J. M. Cherry, A. P. Davis, K. Dolinski, S. S. Dwight, J. T. Eppig, M. A. Harris, D. P. Hill, L. Issel-Tarver, A. Kasarskis, S. Lewis, J. C. Matese, J. E. Richardson, M. Ringwald, G. M. Rubin, and G. Sherlock, Gene Ontology: Tool for the unification of biology, Nature genetics 25, 25 (2000).

[13] K. Zarringhalam, D. Degras, C. Brockel, and D. Ziemek, Robust phenotype prediction from gene expression data using differential shrinkage of co-regulated genes, Scientific Reports 8, 1237 (2018).

[14] K. Zarringhalam, A. Enayetallah, P. Reddy, and D. Ziemek, Robust clinical outcome prediction based on Bayesian analysis of transcriptional profiles and prior causal networks, Bioinformatics 30, i69 (2014).

[15] T. Kang, W. Ding, L. Zhang, D. Ziemek, and K. Zarringhalam, A biological network-based regularized artificial neural network model for robust phenotype prediction from gene expression data, BMC Bioinformatics 18, 565 (2017).

[16] Y.-J. Shen and S.-G. Huang, Improve Survival Prediction Using Principal Components of Gene Expression Data, Genomics, Proteomics & Bioinformatics 4, 110 (2006).

[17] R. Lopez, J. Regier, M. B. Cole, M. I. Jordan, and N. Yosef, Deep generative modeling for single-cell transcriptomics, Nature Methods 15, 1053 (2018).

[18] C. H. Grønbech, M. F. Vording, P. N. Timshel, C. K. Sønderby, T. H. Pers, and O. Winther, scVAE: Variational auto-encoders for single-cell gene expression data, bioRxiv 10.1101/318295 (2018).

[19] G. P. Way and C. S. Greene, Extracting a biologically relevant latent space from cancer transcriptomes with variational autoencoders, Pacific Symposium on Biocomputing. Pacific Symposium on Biocomputing 23, 80 (2018).

[20] L. Rampasek, D. Hidru, P. Smirnov, B. Haibe-Kains, and A. Goldenberg, Dr.VAE: Drug Response Variational Autoencoder, arXiv:1706.08203 [stat] (2017), arXiv:1706.08203 [stat].

[21] Lonsdale et al., The Genotype-Tissue Expression (GTEx) project, Nature Genetics 45, 580 (2013).

[22] Barrett et al., NCBI GEO: Archive for functional genomics data sets–update, Nucleic Acids Re-search 41, D991 (2013).

[23] Liu et al., An Integrated TCGA Pan-Cancer Clinical Data Resource to Drive High-Quality Survival Outcome Analytics, Cell 173, 400 (2018).

[24] G. P. Wagner, K. Kin, and V. J. Lynch, Measurement of mRNA abundance using RNA-seq data: RPKM measure is inconsistent among samples, Theory in Biosciences = Theorie in Den Biowis-senschaften 131, 281 (2012).

[25] B. Li and C. N. Dewey, RSEM: Accurate transcript quantification from RNA-Seq data with or without a reference genome, BMC Bioinformatics 12, 323 (2011).

[26] J. Aitchison, The Statistical Analysis of Compositional Data, Journal of the Royal Statistical Society. Series B (Methodological) 44, 139 (1982).

[27] D. Lovell, V. Pawlowsky-Glahn, J. J. Egozcue, S. Marguerat, and J. Bähler, Proportionality: A valid alternative to correlation for relative data, PLoS computational biology 11, e1004075 (2015).

[28] A. D. Fernandes, J. N. Reid, J. M. Macklaim, T. A. McMurrough, D. R. Edgell, and G. B. Gloor, Unifying the analysis of high-throughput sequencing datasets: Characterizing RNA-seq, 16S rRNA gene sequencing and selective growth experiments by compositional data analysis, Microbiome 2, 15 (2014).

[29] K. Chawla, S. Tripathi, L. Thommesen, A. Lægreid, and M. Kuiper, TFcheckpoint: A curated compendium of specific DNA-binding RNA polymerase II transcription factors, Bioinformatics (Oxford, England) 29, 2519 (2013).

[30] H. B. Mann and D. R. Whitney, On a Test of Whether one of Two Random Variables is Stochastically Larger than the Other, The Annals of Mathematical Statistics 18, 50 (1947).

[31] F. E. Harrell, Regression Modeling Strategies: With Applications to Linear Models, Logistic Regression, and Survival Analysis (Springer Science & Business Media, 2001).

[32] Fabregat et al., The Reactome Pathway Knowledgebase, Nucleic Acids Research 46, D649 (2018).

[33] A. Krämer, J. Green, J. Pollard, and S. Tugendreich, Causal analysis approaches in Ingenuity Pathway Analysis, Bioinformatics (Oxford, England) 30, 523 (2014).

[34] I. V. Ozerov et al. In Silico Pathway Activation Network Decomposition Analysis (iPANDA) as a method for biomarker development, Nature Communications 7, 13427 (2016).

[35] Y. Zhao and R. Simon, Gene expression deconvolution in clinical samples, Genome Medicine 2, 93 (2010).

[36] R. Gaujoux and C. Seoighe, CellMix: A comprehensive toolbox for gene expression deconvolution, Bioinformatics 29, 2211 (2013).

[37] S. S. Shen-Orr, R. Tibshirani, P. Khatri, D. L. Bodian, F. Staedtler, N. M. Perry, T. Hastie, M. M. Sarwal, M. M. Davis, and A. J. Butte, Cell type–specific gene expression differences in complex tissues, Nature Methods 7, 287 (2010).

[38] A. Gupta, H. Wang, and M. Ganapathiraju, Learning structure in gene expression data using deep architectures, with an application to gene clustering, bioRxiv 10.1101/031906 (2015).

[39] A. B. Dincer, S. Celik, N. Hiranuma, and S.-I. Lee, DeepProfile: Deep learning of cancer molecular profiles for precision medicine, bioRxiv 10.1101/278739 (2018).

[40] G. P. Way and C. S. Greene, Evaluating deep variational autoencoders trained on pan-cancer gene expression, arXiv:1711.04828 [q-bio] (2017), arXiv:1711.04828 [q-bio].

[41] C. K. Fisher, A. M. Smith, and J. R. Walsh, Who is this gene and what does it do? A toolkit for munging transcriptomics data in python, bioRxiv, 299107 (2018).

[42] M. Suáarez-Farinñas et al., RNA sequencing atopic dermatitis transcriptome profiling provides insights into novel disease mechanisms with potential therapeutic implications, The Journal of Allergy and Clinical Immunology 135, 1218 (2015).

[43] B. C. E. Peck et al., MicroRNAs Classify Different Disease Behavior Phenotypes of Crohn’s Disease and May Have Prognostic Utility, Inflammatory Bowel Diseases 21, 2178 (2015).

[44] G. W. Tew et al., Association Between Response to Etrolizumab and Expression of Integrin αE and Granzyme A in Colon Biopsies of Patients With Ulcerative Colitis, Gastroenterology 150, 477 (2016).

[45] P. DiMeglio, J. a. H. Duarte, H. Ahlfors, N. D. L. Owens, Y. Li, F. Villanova, I. Tosi, K. Hirota, F. O. Nestle, U. Mrowietz, M. J. Gilchrist, and B. Stockinger, Activation of the aryl hydrocarbon receptor dampens the severity of inflammatory skin conditions, Immunity 40, 989 (2014).

[46] J. a. Fadista et al., Global genomic and transcriptomic analysis of human pancreatic islets reveals novel genes influencing glucose metabolism, Proceedings of the National Academy of Sciences of the United States of America 111, 13924 (2014).

[47] W. R. Swindell, H. A. Remmer, M. K. Sarkar, X. Xing, D. H. Barnes, L. Wolterink, J. J. Voorhees, R. P. Nair, A. Johnston, J. T. Elder, and J. E. Gudjonsson, Proteogenomic analysis of psoriasis reveals discordant and concordant changes in mRNA and protein abundance, Genome Medicine 7, 86 (2015).

[48] R. Arora, A. Cotter, K. Livescu, and N. Srebro, Stochastic optimization for PCA and PLS, in 2012 50th Annual Allerton Conference on Communication, Control, and Computing (Allerton) (2012) pp. 861–868.

[49] P. Vincent, H. Larochelle, I. Lajoie, Y. Bengio, and P.-A. Manzagol, Stacked denoising autoen-coders: Learning useful representations in a deep network with a local denoising criterion, Journal of machine learning research 11, 3371 (2010).

[50] D. P. Kingma and M. Welling, Auto-Encoding Variational Bayes, arXiv:1312.6114 [cs, stat] (2013), arXiv:1312.6114 [cs, stat].

[51] G. Klambauer, T. Unterthiner, A. Mayr, and S. Hochreiter, Self-Normalizing Neural Networks, arXiv:1706.02515 [cs, stat] (2017), arXiv:1706.02515 [cs, stat].

[52] S. R. Bowman, L. Vilnis, O. Vinyals, A. M. Dai, R. Jozefowicz, and S. Bengio, Generating Sentences from a Continuous Space, arXiv:1511.06349 [cs] (2015), arXiv:1511.06349 [cs].

[53] A. Paszke, S. Gross, S. Chintala, G. Chanan, E. Yang, Z. DeVito, Z. Lin, A. Desmaison, L. Antiga, and A. Lerer, Automatic differentiation in PyTorch, (2017).

[54] F. Pedregosa et al., Scikit-learn: Machine Learning in Python, Journal of Machine Learning Research 12, 2825 (2011).

[55] B. Efron, The Efficiency of Cox’s Likelihood Function for Censored Data, Journal of the American Statistical Association 72, 557 (1977).

[56] D. P. Kingma and J. Ba, Adam: A Method for Stochastic Optimization, arXiv:1412.6980 [cs] (2014), arXiv:1412.6980 [cs].

[57] X. Glorot and Y. Bengio, Understanding the difficulty of training deep feedforward neural networks, Proceedings of the thirteenth international conference on artificial intelligence and statistics, 249 (2010).

[58] S. Ioffe and C. Szegedy, Batch Normalization: Accelerating Deep Network Training by Reducing Internal Covariate Shift, arXiv preprint arXiv:1502.03167 (2015).

